# BAX requires VDAC2 to mediate apoptosis and to limit tumor development

**DOI:** 10.1101/266668

**Authors:** Hui San Chin, Mark F. van Delft, Robert L. Ninnis, Mark X. Li, Iris K. L. Tan, Boris Reljic, Kristen Scicluna, Laura F. Dagley, Jarrod J. Sandow, Gemma L. Kelly, Stephane Chappaz, Seong L. Khaw, Catherine Chang, Andrew Webb, Colin Hockings, Cathrine M. Hall, Andrew J. Kueh, Michael T. Ryan, Ruth M. Kluck, Philippe Bouillet, Marco J. Herold, Daniel H. D. Gray, David C. S. Huang, Grant Dewson

**Affiliations:** Walter and Eliza Hall Institute of Medical Research, 1G Royal Parade, Parkville, Melbourne, Victoria 3052, Australia; Department of Medical Biology, University of Melbourne, Parkville, Melbourne, Victoria 3010, Australia; Department of Biochemistry and Molecular Biology, Monash University, Melbourne, Victoria 3800, Australia

**Keywords:** Apoptosis, BAX, BAK, BCL-2, cancer, mitochondria, voltage-dependent anion channel

## Abstract

Intrinsic apoptosis is critical for normal physiology including the prevention of tumor formation. BAX and BAK are essential for mediating this process and for the cytotoxic action of many anticancer drugs. BAX and BAK are thought to act in a functionally redundant manner and are considered to be regulated similarly. From an unbiased genome-wide CRISPR/Cas9 screen, we identified VDAC2 (voltage-dependent anion channel 2) as essential for BAX, but not BAK, to function. The genetic deletion of *VDAC2* abrogated the association of BAX and BAK with mitochondrial complexes that contain VDAC1, VDAC2 and VDAC3. By disrupting its localization to mitochondria, BAX is rendered completely ineffective. Moreover, we defined an interface unique to VDAC2 that is required to drive BAX activity. Consequently, interfering with this interaction or deleting *VDAC2* phenocopied the loss of *BAX*, including impairing the killing of tumor cells by anti-cancer agents such as the BCL-2 inhibitor venetoclax. Furthermore, the ability of BAX to prevent tumor formation was attenuated in the absence of *VDAC2*. Taken together, our studies show for the first time that BAX-mediated apoptosis, but not BAK-mediated apoptosis, is critically dependent on VDAC2, hence revealing the differential regulation of BAX and BAK.

---

Apoptotic cell death is a fundamental process that is essential for embryonic development and immune system homeostasis. BAX and BAK are members of the BCL-2 family of proteins that have essential, but redundant, functions as mediators of intrinsic apoptosis^1, 2, 3^. The activation of BAX and BAK and their consequent self-association permeabilizes the mitochondrial outer membrane (MOM) to instigate cytochrome *c* release and cell death^1^. While BAK is predominantly integrated into the MOM, BAX is predominantly cytosolic. Their distinct subcellular localizations may reflect different rates of “retrotranslocation” from the MOM to the cytosol^4, 5, 6^, although the precise determinants of their recruitment to the MOM to mediate cell killing are unclear. Many chemotherapeutic agents indirectly trigger BAX/BAK-mediated apoptosis whereas BH3-mimetic compounds, such as venetoclax (ABT-199), directly inhibit BCL-2 proteins to drive apoptosis^2, 7, 8^. Venetoclax, which selectively targets BCL-2, has proven highly efficacious for patients with high-risk chronic lymphocytic leukemia (CLL) leading to its approval for treating such patients^9^.

The VDAC channels (VDAC1, VDAC2 and VDAC3) are responsible for the transport of low molecular weight metabolites across the MOM including adenosine triphosphate (ATP) and adenosine diphosphate (ADP). Although genetic evidence has argued against a role for VDAC channels in mediating cytochrome *c* release^10^, VDACs have been proposed to influence apoptosis by interacting with BCL-2 family proteins including BCL-X_L_, BAX and BAK^11, 12, 13, 14^. In this regard, the prevailing dogma is that VDAC2 acts to limit apoptosis by sequestering BAK^15^.

In marked contrast to this, we identified VDAC2 in an unbiased genome-wide screen for factors *required* for BAX to function. In the absence of *VDAC2*, cell killing mediated by BAX, but not BAK, is abolished. Moreover, the interaction with VDAC2 is critical for BAX to mediate cell death in response to chemotherapeutic agents both *in vitro* and *in vivo*, as well as for BAX to limit *in vivo* tumor development. Our genetic and functional studies unequivocally define a critical and unique requirement for VDAC2 in the apoptotic activity of BAX.

## Results

### VDAC2 is required for BAX to mediate apoptosis

To identify novel regulators of apoptosis, we undertook unbiased, genome-wide CRISPR/Cas9 library screens (Fig. 1a). *Mcl1*-deficient mouse embryonic fibroblasts (MEFs) were used as they readily undergo apoptosis when the remaining pro-survival proteins they express (BCL-2, BCL-X_L_, BCL-W) are inhibited by the BH3-mimetic ABT-737 (Supplementary Fig. 1a and b)^16^. *Mcl1*^−/−^ MEFs stably expressing Cas9 were infected with a genome-wide short guide RNA (sgRNA) library (Supplementary Fig. 1c, 87,897 sgRNAs targeting 19,150 mouse genes^17^). Following treatment with ABT-737, surviving cells were harvested and enriched sgRNAs (relative to untreated MEFs) identified by deep sequencing (Fig. 1b). As expected, sgRNAs targeting *Bax* or *Bak* were enriched in *Mcl1*^*−/−*^ MEFs that survived ABT-737 treatment (Fig. 1c, Supplementary Table 1).

**Figure 1.**
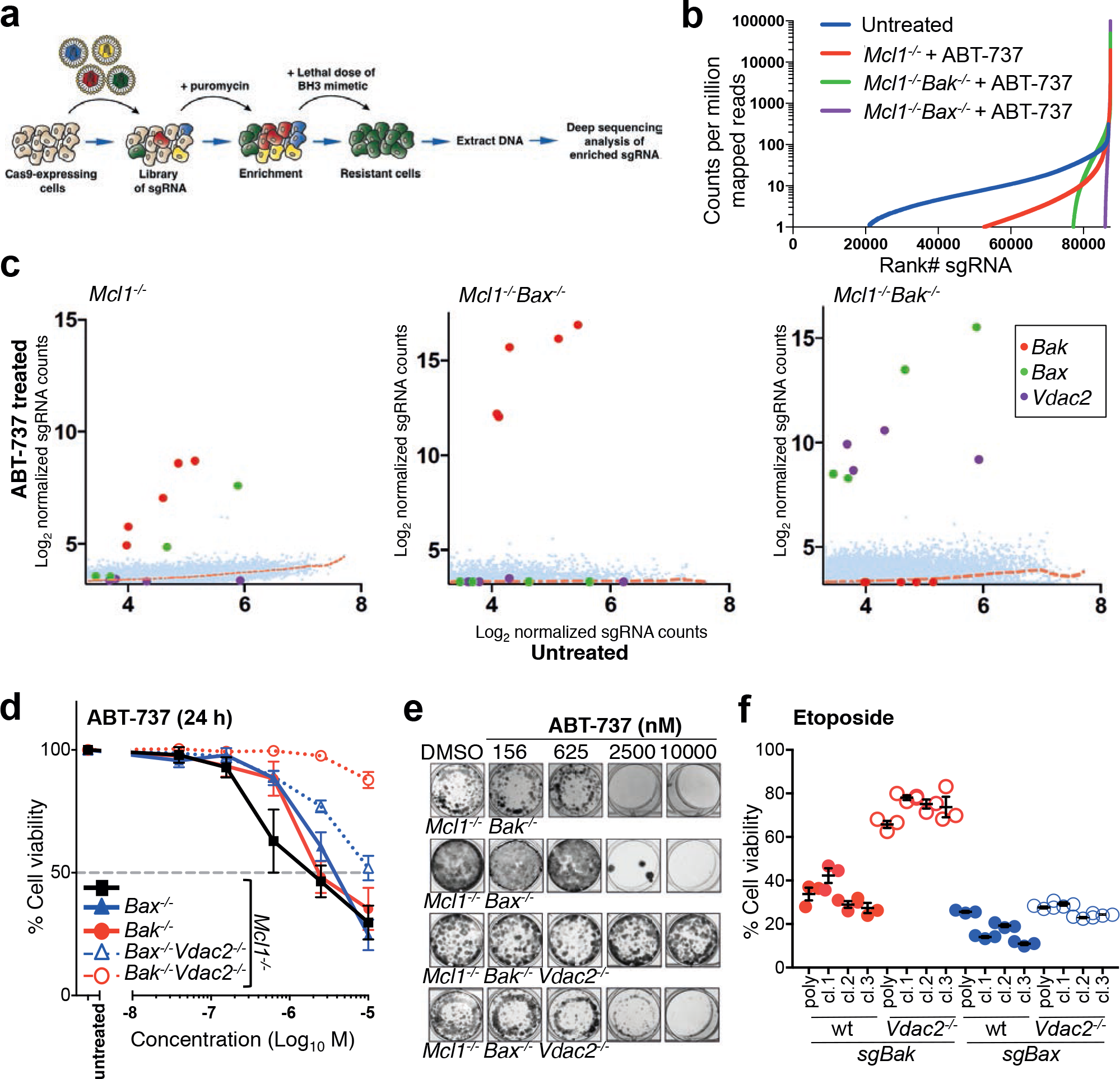
CRISPR/Cas9 whole genome library screen identifies VDAC2 as an essential promoter of BAX-mediated apoptosis. **(a)** Outline of the genome-wide CRISPR/Cas9 library screens to identify mediators of intrinsic apoptosis. **(b)** Reduced sgRNA representation following treatment with ABT-737. In untreated control cells, almost 60,000 sgRNA were detected. Gene ontology analysis confirmed that those sgRNA that were not detected were significantly enriched for essential house-keeping genes (data not shown). Following treatment with 250 nM (for *Mcl1*^*−/−*^) and 350 nM ABT-737 (for *Mcl1*^*−/−*^*Bak*^*−/−*^ and *Mcl1*^*−/−*^ *Bax*^*−/−*^) a reduced pool of sgRNAs was recovered. **(c)** VDAC2 is an essential promoter of BAX apoptotic function. MEFs expressing Cas9 and a whole-genome sgRNA library were treated with ABT-737 (LC90, 250 nM, see Supplementary Fig. 1b) for 48 h. Resistant cells were recovered after 5 days and enriched sgRNAs identified by deep-sequencing. Plots show independent sgRNAs in ABT-737 treated versus untreated controls cells collated and normalized from 2-4 independent experiments. **(d)** Deletion of *Vdac2* protects from BAX-mediated apoptosis in response to ABT-737. Clones (*Mcl1*^*−/−*^*Bax*^*−/−*^*Vdac2*^*−/−*^ and *Mcl1*^*−/−*^*Bak*^*−/−*^*Vdac2*^*−/−*^) or polyclonal populations (*Mcl1*^*−/−*^, *Bax*^*−/−*^ and *Bak* ^*−/−*^) of MEFs were treated with escalating doses of ABT-737 for 24 h and cell viability assessed by PI exclusion. Data is mean +/− SEM of at least 3 independent experiments with 3-4 independent clones. **(e)** Deletion of *Vdac2* provides long term protection from BAX-mediated cell death. MEFs of the indicated genotype were treated with escalating doses of ABT-737 and colony formation was assessed after 5 days. **(f)** Deletion of *Bak* protects *Vdac2*^*−/−*^ MEFs from etoposide-induced apoptosis. Polyclonal populations or 3 independent MEF clones of the indicated genotype (all on 129sv;C57BL/6 background) were treated with etoposide (10 μM for 24 h) and cell viability assessed by PI exclusion. Data is mean +/− SEM shown for 3 independent experiments.

In an attempt to identify factors that may act specifically on BAX or BAK, we undertook sensitizer screens in *Mcl1*^*−/−*^ MEFs lacking either one of the cell death mediators. We undertook the screen in *Mcl1*^*−/−*^ *Bax*^*−/−*^ MEFs, in order to genetically isolate BAK-dependent apoptosis and thus identify genes that may be required for apoptosis mediated by BAK. We failed to identify any such genes, with only sgRNAs targeting *Bak* being over-represented in this screen (Fig. 1c, Supplementary Table 2). Of note, these screens would not have identified any regulators critical for normal cell growth or those acting redundantly to facilitate BAK function. Conversely, when we performed the screen using *Mcl1*^*−/−*^*Bak*^*−/−*^ MEFs to genetically isolate BAX-dependent apoptosis, eight sgRNAs were significantly enriched; four targeting *Bax* and four targeting *Vdac2* (Fig. 1c, Supplementary Table 3). This strongly indicated that deletion of *Vdac2* protected cells from BAX-mediated apoptosis, but not BAK-mediated apoptosis.

To validate the findings of the screen, we deleted *Vdac2* in either *Mcl1*^*−/−*^*Bak*^*−/−*^ or *Mcl1*^*−/−*^*Bax*^*−/−*^ MEFs using CRISPR/Cas9 gene editing (Supplementary Fig. 1d). Strikingly, when killing could only be mediated by BAX, deleting *Vdac2* from multiple independently derived *Mcl1*^−/−^ *Bak*^−/−^ cells made them highly resistant to ABT-737 (Fig. 1d). By contrast, BAK-mediated killing was hardly affected (Fig. 1d). *Mcl1*^*−/−*^ MEFs are also killed by the selective inhibition of BCL-X_L_ with A1331852^18^ (Supplementary Fig. 1e). Deleting *Vdac2* likewise protected *Mcl1*^*−/−*^*Bak*^*−/−*^ cells from this BH3-mimetic (Supplementary Fig. 1e), thus ruling out the possibility that protection was specific to ABT-737 treatment.

Most remarkably, the combined loss of BAK and VDAC2 (but not BAX and VDAC2) also afforded long term protection against ABT-737 (Fig. 1e) and A1331852 (Supplementary Fig. 1f) in clonogenic assays. The critical role for VDAC2 in BAX apoptotic function extends beyond these model systems since deleting *Bak* in *Vdac2*^*−/−*^MEFs likewise rendered these cells highly resistant to etoposide-induced apoptosis (Fig. 1f). Collectively, our data strongly indicate that apoptosis mediated by BAX, but not BAK, requires VDAC2. Next, we asked whether VDAC2 physically interacts with BAX to promote its activity.

### VDAC2 and VDAC3 are the key components of a BAX-containing mitochondrial complex

We, and others, have reported that BAX and BAK reside in large VDAC2-containing mitochondrial complexes, but become dissociated following the induction of apoptosis^12, 13, 19^. To determine whether these are identical or distinct VDAC2 complexes we performed antibody gel-shift assay on mitochondrial fractions prepared from HeLa or HCT-116 cells (Fig. 2a). We found that adding Fab fragments of an antibody that binds inactive human BAK (7D10^20^) altered the mobility of the BAK:VDAC2 complex on a native gel. However, the BAX:VDAC2 complex was unaffected (Fig. 2a). Hence, VDAC2 must form distinct complexes with BAX or BAK.

**Figure 2.**
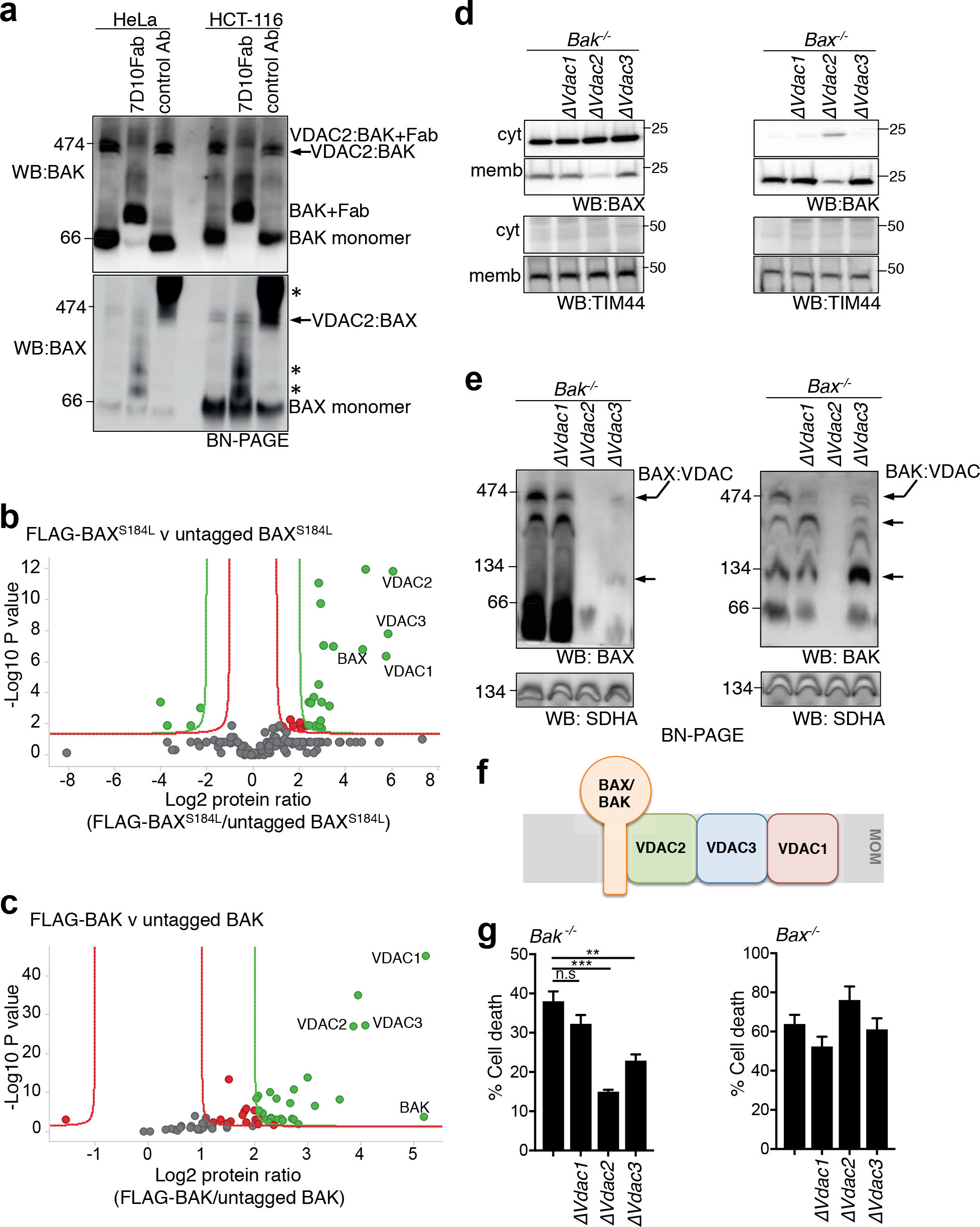
VDAC2 promotes the stable association of BAX and BAK with a mitochondrial complex involving VDAC1, VDAC2 and VDAC3. **(a)** Endogenous BAX and BAK associate with independent complexes in mitochondria. Mitochondria-enriched fractions from HeLa or HCT-116 cells were incubated with a control antibody or an antibody Fab fragment that binds inactive human BAK (7D10), prior to BN-PAGE and immunoblotting for BAK or BAX. * indicates cross-reactivity of anti-rat secondary antibody with rat IgG used for gel-shift. **(b)** Mass spectrometry of the native BAX complex. Mitochondria from MEFs expressing FLAG-BAX^S184L^ or untagged BAX^S184L^ were solubilized in 1% digitonin prior to anti-FLAG affinity purification and proteins identified by mass spectrometry. **(c)** Mass spectrometry of the native BAK complex. Mitochondria from MEFs expressing FLAG-BAK or untagged BAK were solubilized in 1% digitonin prior to anti-FLAG affinity purification and proteins identified by mass spectrometry. **(d)** Deletion of VDAC2 impacts mitochondrial localization of BAX and BAK. Clonal populations of *Bax*^*−/−*^ and *Bak*^*−/−*^ MEFs with deleted *Vdac1, Vdac2* or *Vdac3* (denoted Δ) were fractionated into cytosol and membrane and immunoblotted for BAX, BAK or TIM44 as a mitochondrial control. **(e)** VDAC2 plays the major role in BAX and BAK complex stability. Mitochondria isolated from clonal populations of *Bax*^*−/−*^ and *Bak*^*−/−*^ MEFs with deleted *Vdac1, Vdac2* or *Vdac3* were analyzed on BN-PAGE. Data is representative of two independent clones (see Supplementary Fig. 2e). Intermediate complexes indicated (arrows). **(f)** Proposed functional hierarchy and topology of the BAX/BAK:VDAC complex based on biochemical and functional analysis. **(g)** BAX-mediated apoptosis is impaired in the absence of VDAC2 and to a lesser extent by VDAC3. Polyclonal populations were treated with etoposide (10 μM) and cell death was assessed by PI uptake. Data is mean+/-SEM of 3 independent experiments. ***p<0.001; **p<0.01; n.s, not significant; based on unpaired Student’s t-test.

The sizes of these complexes on native gels also suggested that they are large multimeric complexes that could contain other components. To identify constituent proteins, we generated MEFs stably expressing FLAG-BAX^S184L^ or FLAG-BAK (both of which are constitutively localized to mitochondria through an interaction with VDAC2^13^) and purified the complexes under native conditions using an anti-FLAG antibody (Supplementary Fig. 2a). While mass spectrometry of these complexes confirmed the presence of VDAC2 and BAX or BAK, VDAC1 and VDAC3 were also present (Fig. 2b and c, Supplementary Tables 4 and Table 5).

In order to define the role of individual VDACs in BAX or BAK complex formation, we next evaluated the impact of deleting each VDAC (Supplementary Figs. 2b and 2c) on the subcellular localization of BAX and BAK (Fig. 2d) and mitochondrial complex formation (Fig. 2e, and Supplementary Figs. 2d and 2e). Deleting *Vdac2* had a profound effect on BAX: the localization of BAX to mitochondria was disrupted (Fig. 2d) and mitochondrial VDAC2:BAX complexes dissipated (Fig. 2e and Supplementary Fig. 2e). While deleting *Vdac3* also impacted on BAX complex formation (Fig. 2e and Supplementary Fig. 2e), mitochondrial localization of BAX was unaffected (Fig. 2d). Likewise, VDAC2 played an important role in the mitochondrial localization of BAK (Fig. 2d), whilst VDAC3 played an ancillary role indicated by the modified mitochondrial complexes following *Vdac3* deletion (Fig. 2e).

These biochemical studies suggest that the constitution of the distinct mitochondrial VDAC:BAX or VDAC:BAK complexes are largely similar (Fig. 2b and 2c). VDAC2 appeared critical for the formation of both BAX and BAK complexes (Fig. 2e and Supplementary Fig. 2e) and for their localization on mitochondria (Fig. 2d). That VDAC3 played an ancillary role (Fig. 2d, 2e and Supplementary Fig. 2e), suggests a functional hierarchy between the VDACs and implies a potential topology of the mitochondrial complexes (Fig. 2f). Consistent with this notion, we found that VDAC2 had the greatest role in BAX-mediated apoptosis, with VDAC3 playing a lesser role, and VDAC1 largely dispensable (Fig. 2g). This clear functional distinction between VDAC2 and VDAC1 for BAX to function allowed us to clarify how VDAC2 might promote the activity of BAX.

### A distinct surface on VDAC2 drives BAX activity

Of note, re-expression of VDAC2, but not VDAC1, rescued BAX complex formation in VDAC2-deficient cells (Fig. 3a). Moreover, while VDAC2 could rescue BAX driven apoptosis, VDAC1 could not (Fig. 3b). We exploited this distinctive functional impact when these closely related VDACs were expressed to map the precise region on VDAC2 necessary to promote BAX activation. To do this, we generated and expressed VDAC1/2 chimeras (Fig. 3c and Supplementary Fig. 2f). We identified a region of VDAC2, comprising central β-strands 7-10 that was sufficient to promote BAX apoptotic function (Fig. 3d), as well as support the formation of BAX complexes in the mitochondrial membrane (Fig. 3e).

**Figure 3.**
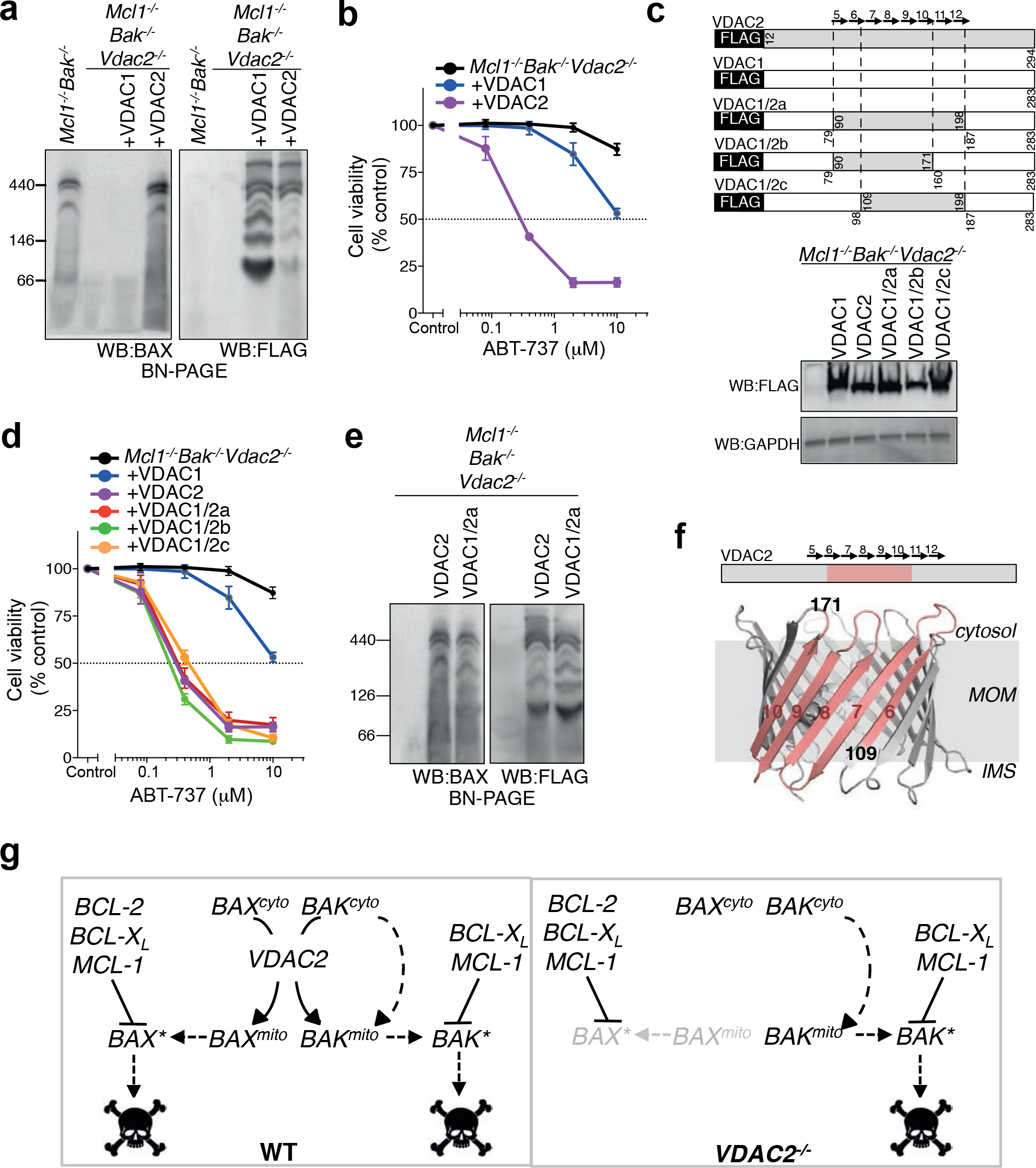
Interaction with VDAC2 is important for BAX apoptotic function. **(a)** BAX mitochondrial complex formation specifically relies on VDAC2. Mitochondria-enriched fractions from *Mcl1*^*−/−*^*Bak*^*−/−*^*Vdac2*^*−/−*^ MEFs reconstituted with FLAG-mVDAC1 or FLAG-hVDAC2 were analyzed by BN-PAGE and immunoblotted for BAX (*left*) or FLAG to detect ectopically-expressed VDACs (*right*). **(b)** BAX apoptotic function relies on VDAC2. Cells as in (a) were treated with ABT-737 and cell viability was assessed. **(c-e)** Rescue of BAX apoptotic function correlates with interaction with a specific region of VDAC2. *Mcl1*^*−/−*^*Bak*^*−/−*^*Vdac2*^*−/−*^ MEFs stably expressing FLAG-mVDAC1/hVDAC2 chimeras (c) were analyzed for expression by immunoblotting for FLAG (or GAPDH as a loading control), cell viability following treatment with ABT-737 (d), and complex formation by BN-PAGE and immunblotting for BAX or FLAG (e). **(f)** BAX interacts with a defined region of VDAC2. Interaction of BAX requires aa109-171 of hVDAC2 (salmon) mapped onto the structure of zebrafish VDAC2 (PDB 4BUM^57^). MOM, mitochondrial outer membrane; IMS, intermembrane space. **(g)** VDAC2 is essential for BAX, but not BAK to target mitochondria and mediate apoptosis. In wild-type cells, cytosolic BAX (BAX^cyto^) relies on VDAC2 to associate with mitochondria (BAX^mito^) and activate (BAX*). In the absence of VDAC2, BAX cannot mediate cell death. Although the ability of BAK to associate with mitochondria is also perturbed in *Vdac2*^*−/−*^ cells^13, 29^, sufficient BAK can still target mitochondria to mediate apoptosis presumably by a VDAC2-independent mechanism. Data presented in (b) and (d) is mean +/− SEM of 3 independent experiments.

Taken together, we postulate that a distinct surface on VDAC2 is required for the recruitment of BAX to the mitochondria and that disrupting this completely abrogates BAX function (Figs. 3f and 3g). Interestingly, the same region of VDAC2 appears to mediate its interaction with BAK^14^, suggesting a conserved mechanism for recruiting both BAX and BAK to mitochondria. However, the functional impact of *Vdac2* deletion on the apoptotic function of BAX and BAK differs. That BAK still drives apoptosis without VDAC2 (Figs. 1c, 1d and 2g), indicates that VDAC2 is not the sole conduit for BAK to the MOM where it acts (Fig. 3g). Conversely, VDAC2 is essential for BAX recruitment to mitochondria and its absence nullifies BAX apoptotic function (Fig. 3g).

### BAK does not limit embryonic development of Vdac2^−/−^ mice

Contrary to our studies uncovering the essential role of VDAC2 in promoting BAX-mediated apoptosis, it had been reported that VDAC2 inhibits BAK^15^. Hence, it was proposed that the early embryonic lethality upon deleting *Vdac2* is caused by inappropriate and excessive apoptosis driven by unrestrained BAK^15^. If so, we should expect that co-deleting *Bak* should allow Vdac2-null embryos to survive. To test this hypothesis, we injected C57BL/6 zygotes with plasmid encoding Cas9 together with short guide RNAs targeting *Bak* and *Vdac2* (Fig. 4a). The resulting embryos were harvested at E14.5 and although all appeared grossly normal (data not shown), genotyping of these embryos was revealing. Most embryos were *Vdac2*-null (87.5%), yet concomitant *Bak* deletion was only observed in 8% compared with 23% when *Bak* was targeted alone (Fig. 4a). This suggested that co-deletion of *Bak* neither promoted nor was it essential for the survival of *Vdac2*^*−/−*^ embryos, at least up to the E14.5 developmental stage.

**Figure 4.**
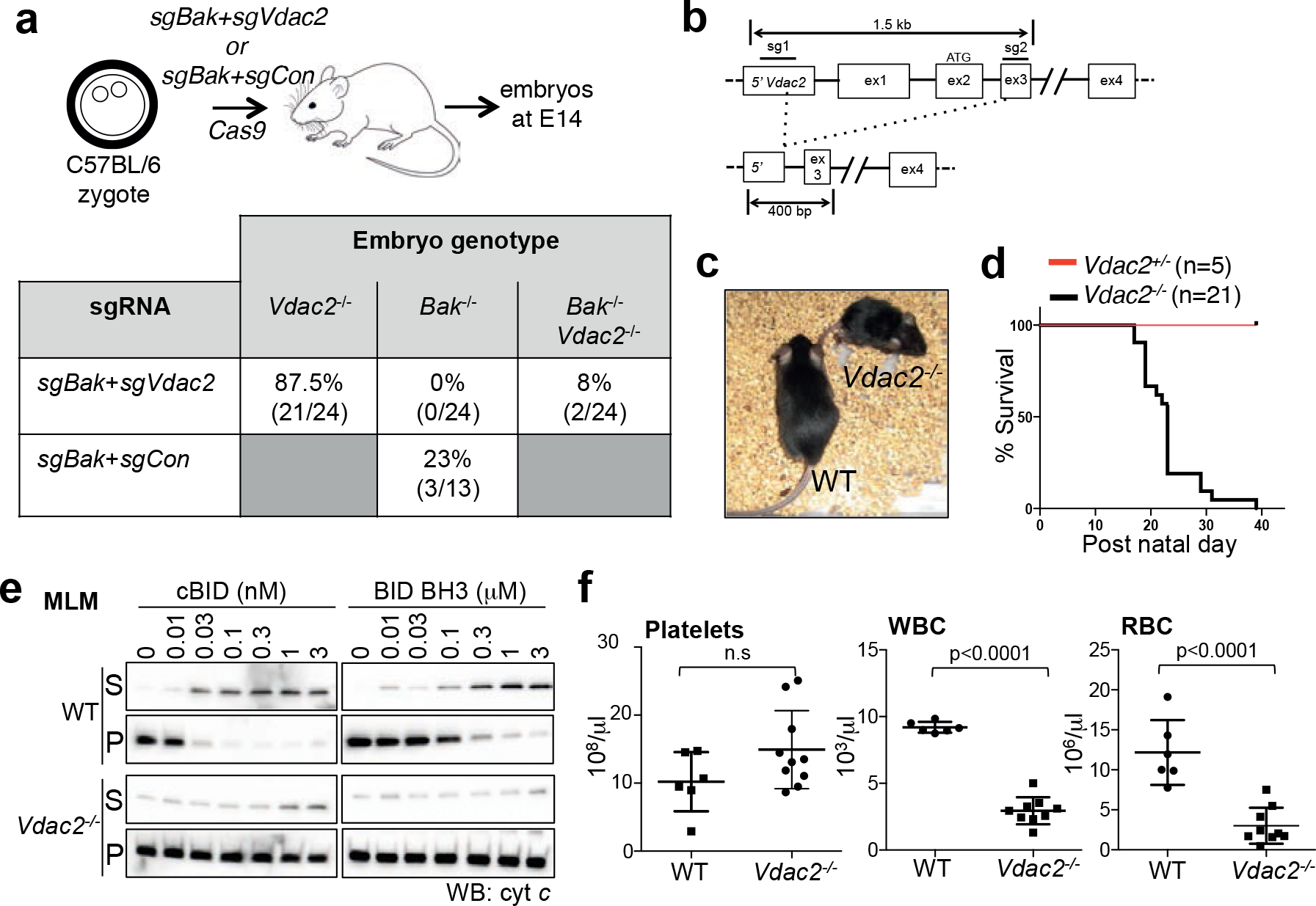
*Vdac2*^*−/−*^ mice do not show evidence of excessive BAK activity. **(a)** *Bak* deletion is not essential for early embryonic development of *Vdac2*^*−/−*^ mice. C57BL/6 zygotes were injected with DNA encoding Cas9 and sgRNAs targeting *Bak* and *Vdac2* (or *Rosa* as a control) and transplanted into pseudo-pregnant mothers. The percentage embryos at E14.5 with homozygous null alleles are indicated (number of mice in parentheses). **(b)** Gene targeting strategy to generate *Vdac2*^*−/−*^ mice. Targeting by both sgRNAs will result in a deletion of 500 bp whereas targeting by the 3 sgRNA alone results in indels (see Supplementary Fig. 3a). **(c and d)** *Vdac2*^*−/−*^ mice are runted die post-natally. Kaplan-Meier survival curve of *Vdac2*^*+/−*^ and *Vdac2*^*−/−*^ F0 mice. **(e)** *Vdac2*^*−/−*^ mitochondria are resistant to MOM permeabilization. Liver mitochondria isolated from age-matched WT and *Vdac2*^*−/−*^ mice were treated with cBID prior to fractionation into supernatant (S) and membrane (P) and immunoblotting for cytochrome *c*. Data are representative of N=2 mice (see Supplementary Fig. 3d). **(f)** *Vdac2*^*−/−*^ mice show defects in the hematopoietic system. Blood counts for individual age-matched wild-type (WT, N=6) or *Vdac2*^*−/−*^ (N=9) mice shown with mean +/−SD. WBC, white blood cells; RBC, red blood cells. P values calculated by twotailed Student’s t-test. n.s, not significant.

Since a significant number of *Vdac2*^*−/−*^ embryos survived to E14.5, we set out to determine whether its loss might compromise development later during embryogenesis. *Vdac2*^*−/−*^ mice were generated on a C57BL/6 genetic background and we allowed the mice to develop to term (Fig. 4b). Of the 26 surviving offspring, 20 were *Vdac2* knock-out with homozygous non-sense (*indels*) mutations (Supplementary Fig. 3a). The absence of VDAC2 protein was confirmed by mass spectrometry and immunoblotting of liver extracts from *Vdac2*^*−/−*^ mice (Supplementary Fig. 3b). Thus, the early embryonic lethality associated with deleting *Vdac2* must depend on genetic background, being much more severe in 129Sv;C57BL/6 mice^15^ compared to the inbred C57BL/6 used here.

### Vdac2^−/−^ mice do not exhibit evidence of excessive BAK-mediated apoptosis

Although *Vdac2*^*−/−*^ mice were viable at birth, they failed to gain weight and had to be euthanized by 6 weeks of age because of ill health (Fig. 4c, 4d and Supplementary Fig. 3c). Regardless, this provided us with an opportunity to investigate the interaction between VDAC2 and BAK *in vivo* in addition to that during embryogenesis, focusing initially on testing if VDAC2 acts to restrain BAK ^15^. Firstly, it is well recognized that BAK-mediates cytochrome *c* release, indicative of mitochondrial outer membrane damage, when mouse liver mitochondria (MLM) are treated with BH3 peptides such as cBID^21^. Surprisingly, this was not enhanced by the deletion of *Vdac2* (Fig. 4e and Supplementary Fig. 3d), but was instead compromised, most probably because BAK levels are reduced in *Vdac2*-null cells (Supplementary Figs. 3e and 3f)^13, 22^.

Secondly, platelet homeostasis is highly dependent on BAK; its deletion causes elevated platelet counts whereas unrestrained BAK leads to low platelet counts^23^. If VDAC2 acts to restrain BAK by their association in platelets (Supplementary Fig. 4a), then deleting VDAC2 should decrease platelet counts, but this was not the case (Fig. 4f). Collectively, our data (Figs. 4a, 4e and 4f) suggest that BAK at mitochondria is not hyperactive in primary tissues that lack VDAC2 and its activation is unlikely to account for the premature demise of *Vdac2*^*−/−*^ mice.

Consistent with this, we found that *Vdac2*-deficient hematopoietic stem cells (HSCs) could function normally (Supplementary Fig. 4b), and apoptosis of thymocytes and splenocytes in *ex vivo* cultures (which can be mediated by either BAX or BAK^24^) was similar in mice reconstituted with wild-type or *Vdac2*^*−/−*^ progenitors (Supplementary Fig. 4c). This suggested that the overall reduction in white and red blood cell counts in *Vdac2*^*−/−*^ mice (Fig. 4f) is secondary to their overall ill health rather than due to excessive BAK-mediated apoptosis. Further investigations revealed the likely cause of their early death. *Vdac2*^*−/−*^ mice had pallid livers (Supplementary Fig. 4d) and their hepatocytes were swollen with central nuclei and clear, distended cytoplasm indicative of cellular edema (Supplementary Fig. 4e). Hydropic swelling often reflects a loss of ionic homeostasis caused by defects in plasma membrane ATP-dependent Na^+^/K^+^ exchange, which can be a consequence of defective mitochondrial ATP production^25^. This liver phenotype would be consistent with the loss of VDAC2 as a transporter of ATP across the MOM, and probably explains why *Vdac2*^*−/−*^ mice fail to thrive.

Although *Vdac2*^*−/−*^ mice showed no signs of hyperactive BAK, importantly, the only VDAC2-deficient male that lived long enough (5 weeks) for such an examination had small testes devoid of mature sperm with expanded numbers of spermatogonia and the appearance of giant cells in seminiferous tubules (Supplementary Fig. 4f). Interestingly, this is a phenotype reminiscent of that observed in *Bax*-deficient mice^26^. This finding is consistent with the notion that VDAC2 promotes the apoptotic function of BAX.

### VDAC2 is required for BAX to mediate tumor cell killing by chemotherapy

Given that our data unequivocally implicates a central role for VDAC2 in promoting BAX-mediated apoptosis, we hypothesized that the BAX:VDAC2 interaction would also be important for the action of BAX in response to chemotherapeutic agents. Apoptosis of HCT116 colorectal cancer cells in response to either ABT-737^27^ or the BCL-X_L_ inhibitor A1331852 relies on BAX (Fig. 5a). Deletion of *VDAC2* rendered HCT116 cells as resistant to these BH3-mimetic compounds as the loss of BAX alone and remarkably, even the combined loss of BAX and BAK (Fig. 5a and Supplementary Fig. 5a). When apoptosis in the same cells could be also driven by BAK (e.g. combining ABT-737 with actinomycin D treatment), the loss of VDAC2 alone had no impact (Fig. 5a).

**Figure 5.**
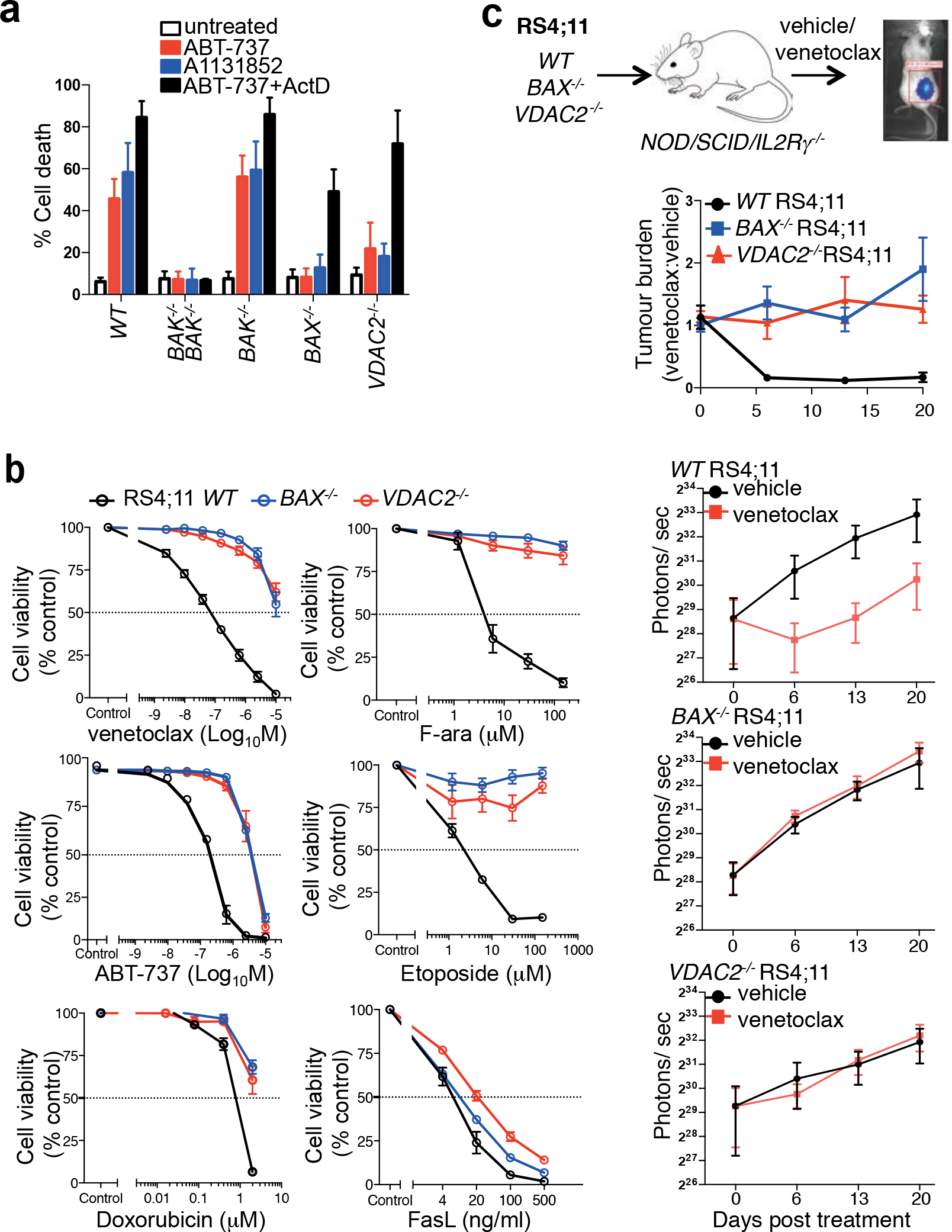
BAX-mediated killing of cancer cells *in vitro* and *in vivo* is dependent on VDAC2. **(a)** Deletion of *VDAC2* protects human cells from apoptosis. HCT116 cells were treated with ABT-737 (5 μM), A1331852 (5 μM) or ABT-737 (5 μM) + actinomycin D (1 μM) for 24 h and cell death assessed. Data is mean +/-SEM of 5 independent experiments. **(b)** BAX or VDAC2 deletion renders RS4;11 acute lymphoblastic leukemia cells resistant to venetoclax or other chemotherapeutic agents. WT, *Bax*^*−/−*^ or *Vdac2*^*−/−*^ RS4;11 cells were treated with venetoclax, ABT-737 and standard-of-care chemotherapies (F-ara, etoposide, doxorubicin) or the BAX/BAK-independent stimulus Fas ligand (FasL) and cell viability assessed by PI exclusion. Data is mean +/−SEM of at least 3 independent experiments. **(c)** Deletion of *VDAC2* renders RS4;11 cells resistant to venetoclax *in vivo*. RS4;11 cells were subcutaneously engrafted into NOD/SCID/IL-2Ryγ^null^ mice, treated with venetoclax and tumor growth monitored by IVIS imaging. The collated data (*top panel*) is normalized to represent relative tumor burden. Data is mean +/− SEM collated from 4 independent experiments, N=12 mice engrafted with each genotype of RS4;11 cells.

Similarly, killing of the acute lymphoblastic leukemia cell line RS4;11 induced by the BCL-2 inhibitor venetoclax (also known as ABT-199)^7, 9^, or standard-of-care chemotherapeutic agents is principally mediated by BAX (Fig. 5b). Independent RS4;11 clones with targeted deletion of *VDAC2* were as resistant to these drugs as *BAX*^*−/−*^ RS4;11 cells (Fig. 5b and Supplementary Fig. 5b). As expected, BAX- or VDAC2-deficient leukemia cells demonstrated comparable sensitivity to activation of the extrinsic pathway of apoptosis by Fas ligand (Fig. 5b)^28^. To test whether the BAX:VDAC2 axis contributes to the response to BH3-mimetics *in vivo*, RS4;11 cells expressing a luciferase reporter were engrafted subcutaneously into NOD/SCID/IL-2Rγ^null^ mice. Deletion of either BAX or VDAC2 was sufficient to cause marked resistance to *in vivo* venetoclax treatment (Fig. 5c).

### VDAC2 is required for BAX to limit tumor formation

As the intrinsic apoptotic pathway is an important barrier to tumorigenesis, we hypothesized that the impairment of BAX-mediated apoptosis by deletion of *Vdac2*, when combined with *Bak* deletion, should accelerate tumor development. Fetal liver-derived HSCs derived from *Bak*^*−/−*^, *Bak*^*−/−*^*Bax*^*−/−*^ or *Bak*^*−/−*^*Vdac2*^*−/−*^ mice at E14.5 were retrovirally transduced with the *c-MYC* oncogene and transplanted into lethally-irradiated mice to induce acute myeloid leukemia (AML) (Fig. 6 and Supplementary Fig. 6)^29^. As expected, the combined loss of *Bax* and *Bak* in hematopoietic progenitors accelerated the development of MYC-driven AML compared with loss of *Bak* alone (Fig. 6). Strikingly, the combined deletion of *Vdac2* and *Bak* also accelerated AML development when compared with deletion of *Bak* alone (Fig. 6), affirming that VDAC2 is a key driver of BAX activation in the context of oncogenic stress.

**Figure 6.**
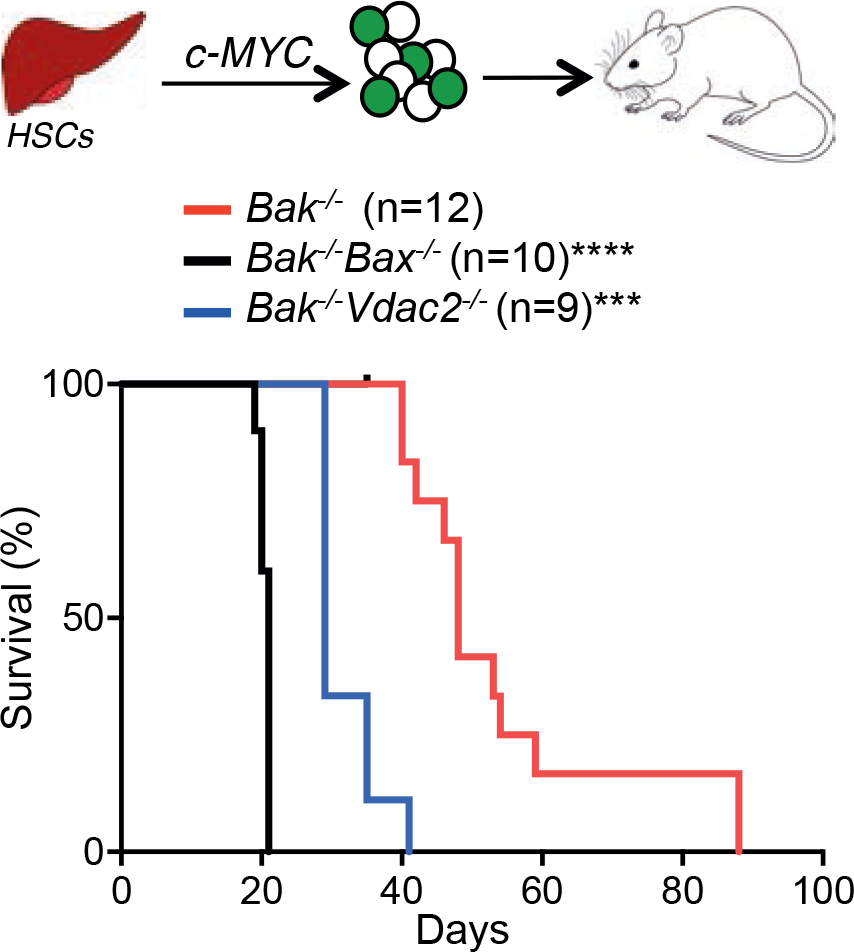
The ability of *Bax* to limit tumor development relies on *Vdac2*. *Vdac2* deletion accelerates the development of *MYC*-driven AML. Kaplan-Meier survival plot of mice transplanted with *Bak*^*−/−*^, *Bak*^*−/−*^*Bax*^*−/−*^ or *Bak*^*−/−*^*Vdac2*^*−/−*^ fetal liver-derived hematopoietic stem cells (HSCs) (2 livers per genotype) infected with a *c-MYC*-expressing retrovirus. P values (Log-rank analysis) of mice injected with *Bak*^*−/−*^ hematopoietic precursors compared with *Bak*^*−/−*^*Vdac2*^*−/−*^ or *Bak*^*−/−*^*Bax*^*−/−*^ precursors are <0.001 (***) and <0.0001 (****) respectively.

## Discussion

Detailed understanding of how intrinsic apoptosis is controlled has paved the way for the development and clinical success of small molecule agonists of the pathway, such as the BCL-2 inhibitor, venetoclax (ABT-199) to treat certain cancers^7, 9, 30^. BAX and BAK act in a functionally redundant manner to mediate intrinsic apoptosis triggered by venetoclax and also during normal tissue homeostasis. Hence, how these essential cell death mediators are regulated is critical both for understanding normal cell death control and for our efforts to target this pathway for therapeutic benefit.

In this regard, VDAC2 plays an intriguing and controversial role. A previous report suggested that VDAC2 restrains BAK and that the early embryonic lethality observed in *Vdac2*-deficient mice is potentially due to unrestrained BAK activity^15^. Unlike the mice on a mixed genetic background used in that study, we found instead that *Vdac2*-null mice on an inbred C57BL/6 genetic background were viable at birth, although they failed to thrive postnatally (Fig. 4C and D). The most prominent pathological finding in these *Vdac2*^*−/−*^ mice was pallid livers filled with swollen hepatocytes. These edematous cells were likely to have resulted from the loss of normal ionic balance across the plasma membrane by ATP-dependent Na^+^/K^+^ pumps^31^, ^32^. These findings suggest that the metabolite transporter function of VDAC2 is important for hepatocyte homeostasis and proper liver function. That *Vdac1*^*−/−*^ and *Vdac3*^*−/−*^ mice are viable and do not exhibit such a liver phenotype or premature decline, implies a non-redundant isoform-specific function for VDAC2^15, 33, 34^. In the context of our study, we failed to find any evidence of heightened sensitivity to apoptosis in various cell types, which would have been expected if BAK was left unrestrained by the loss of VDAC2. Definitive conclusions about the role of VDAC2 in controlling apoptosis *in vivo* are complicated by its dual functions in metabolite transport and in apoptosis and the significant impact of *Vdac2* deletion on the overall health of knockout mice. However, our studies suggest that the loss of *Vdac2* does not promote BAK-mediated apoptosis: 1) platelet number was not reduced (Fig. 4f); 2) liver mitochondria were not more sensitive to cytochrome *c* release when treated with cBID (Fig. 4e); 3) there was no increase in blood cell apoptosis *ex vivo* (Supplementary Fig. 4c).

In parallel studies and in accord with other reports^13, 19, 22^, we confirmed that BAX and BAK interact with VDAC2 on the MOM through a conserved mechanism (Fig. 3). However, the sharply contrasting consequences of blocking VDAC2-driven mitochondrial recruitment on the apoptotic activity of BAX or BAK was completely unanticipated. Using a range of *in vitro* and *in vivo* systems, we discovered that this step is crucial specifically for BAX, but not BAK, to mediate apoptosis. Thus, where BAX is the key mediator of apoptotic cell death, we found identical consequences when either *BAX* or *VDAC2* were genetically deleted. For example, BAX-mediated killing of cancer cells by venetoclax was completely abrogated in the absence of VDAC2. Consistent with these observations we noted defective spermatogenesis upon deleting *Vdac2*, a phenotype reminiscent of *Bax* loss^26^. Based on our studies, we conclude that VDAC2 is essential for recruiting BAX to mitochondria.

Given the proposed redundancy between BAX and BAK, it is somewhat surprising that deleting VDAC2 did not impair BAK activity. This may be because, although VDAC2 plays a role in the localization of BAK to mitochondria^13^, it may not be solely responsible. Properties inherent to BAK itself, such as its C-terminal tail or other factors on the MOM could act independently of VDAC2 to promote BAK localization to mitochondria and hence its apoptotic activity (Fig. 3g). Intriguingly, the removal of VDAC2 renders BAK hyper-active in some situations^13, 15^. A possible explanation is that both BAX and BAK need to dissociate from VDAC2 to homo-oligomerize to mediate cytochrome *c* release^12, 13^. As the residual population of BAK that targets mitochondria independent of VDAC2 does not have to undergo this rate-limiting step, it can directly recruit cytosolic BAK to efficiently mediate MOM permeabilization^13^.

Taken together, we have shown that the VDAC2:BAX interaction is essential for BAX to mediate apoptosis and this clearly differentiates it from BAK. Thus, we hypothesize that manipulating this interaction may well be therapeutically beneficial in situations where BAX, rather than BAK, is the principal mediator of apoptosis. For example, disrupting or preventing the VDAC2:BAX interaction may be a novel strategy to block BAX activity and hence protect cells in some scenarios from damaging cell death. Disrupting the VDAC2:BAX interaction could be exploited to limit pathological apoptosis following traumatic or ischemic brain injuries since differentiated neurons lack functional BAK^35^. In the context of cancer chemotherapy, BAX is likely the prime driver for certain cytotoxic agents to act (Fig. 5)^27, 36^. Our findings are also highly relevant for treatment with venetoclax since its target, BCL-2, principally limits BAX rather than BAK^37^. Venetoclax is now approved as monotherapy for patients with poor-risk chronic lymphocytic leukemia in the US and an increasing number of other countries, but not all patients respond upfront and in some others, secondary resistance develops^9, 38^. Thus, our findings suggest that we should also screen for silencing of VDAC2, in addition to BAX^36^, as a potential driver of resistance to venetoclax.

## Acknowledgments

Jason Corbin for Advia blood analysis, Crystal Stilvala, Nicole Lynch and Shannon Oliver for animal husbandry, Reema Jain, Antonia Policheni and Wenxi Zhou for assistance. We thank Andreas Strasser, Benjamin Kile, Marc Pellegrini, and Zina Valaydon for critical comments and advice, Guillaume Lessene and Jean-Marc Garnier for A1331852, and William Craigen for WT and *Vdac2*^*−/−*^ MEFs. This work was supported by scholarships to HSC from Melbourne University (MIRS, MIFRS) and the Walter and Eliza Hall Institute of Medical Research (Edith Moffatt), fellowships (1043149 to DCSH and 1090236 to DHDG) and grants (1016701, 1057742, 1059290, 1078763, 1078924, 1083077, 1113133) from the Australian National Health and Medical Research Council, a Leukaemia & Lymphoma Society (SCOR grant 7001-13), the Australian Cancer Research Foundation, a fellowship from the Australian Research Council (FT100100791 to GD), the Cancer Council Victoria (grants-in-aid to DCSH and DHDG) and operational infrastructure grants through the Australian Government IRISS and the Victorian State Government OIS 9000220.

## Author contributions

DCSH, GD, MFvD conceived the study; DCSH, DHDG, GD, HSC, MFvD, designed experiments; CC, CMH, CH, DHDG, GD, GLK, HSC, IKLT, RLD, LFD, AW, JJS, KS, MFvD, SC, SLK, XL performed experiments and interpreted data; AJK, MJH generated gene targeted mice; BR, MTR, PB, RMK provided critical advice and reagents; DCSH, GD, HSC, MFvD wrote the manuscript; all authors contributed to preparing the manuscript.

## Materials & Correspondence

Correspondence and request for materials should be addressed to G. Dewson.

## Materials & Methods

### Animal models

All mice were in an inbred C57BL/6J genetic background. All animal experiments conformed to the regulatory standards of, and were approved by, the Melbourne Health Research Directorate Animal Ethics Committee.

### Isolation of mouse liver mitochondria

The isolation of mitochondria from the livers of 4-6 week old mice was performed as described^39, 40^. Mitochondria were incubated with recombinant caspase-8 cleaved BID (cBID) or a BID BH3 peptide (DIIRNIARHLAQVGDSMDRSIPPG) and incubated at 37°C for 2h prior to separation of soluble and membrane fractions and immunoblotting for cytochrome *c* and BAK.

### Cell culture, retroviral infection, CRISPR/Cas9 gene targeting of cell lines, cell survival assays

Mouse embryonic fibroblasts (MEFs) isolated from embryos at embryonic day 14.5 were transformed with SV40-large T antigen as described^41^. MEFs were passaged in Dulbecco’s Modified Eagles Medium supplemented with 10% fetal calf serum (FCS), 55 μM 2-mercaptoethanol and 250 μM asparagine. HeLa, HCT116 colorectal cancer cells and RS4;11 acute lymphoblastic leukemia cells were passaged in RPMI supplemented with 10% FCS. Cells were cultured at 37°C and 10% CO_2_. The wild-type and *Vdac2*^*−/−*^ MEFs used in Fig. 1f were derived from 129Sv;C57BL/6 mice, otherwise all MEFs were derived from C57BL/6 inbred mice.

FLAG-VDAC constructs in the vector pMX-IRES-hygromycin were retrovirally transduced into MEFs as previously described^42^, and transduced cells were selected with hygromycin.

For CRISPR/Cas9 gene editing, parental cells were lentivirally infected with constructs encoding Cas9 (*mCherry*) and doxycycline-inducible short guide RNA (*GFP*) targeting early protein coding exons of the desired gene designed using CRISPR design software (crispr.mit.edu)^43^. Following selection of transduced cells by sorting double positive cells on a FACS AriaII flow cytometer (Becton Dickinson) (Supplementary Fig. 1c), cells were treated with 1 μg/mL doxycycline hyclate (Sigma) to induce sgRNA expression. Alternatively, cells were transiently transfected with sgRNA cloned into PX458 (a kind gift from Feng Zhang^44^) and individual clones were selected. Gene targeting of polyclonal populations (denoted ‘Δ’) was confirmed by immunoblotting or of independent clones (denoted ‘^−/−^’) by next generation sequencing. HCT116 *Vdac2*^*−/−*^ cells were generated by TALEN gene editing as previously reported^13^.

To test apoptotic response, cells were treated with venetoclax, ABT-737, ABT-263, etoposide, fludarabine or FasL. Short-term cell death response was assessed by propidium iodide uptake and flow cytometry using a FACS Calibur flow cytometer (Becton Dickinson). Longterm clonogenic potential was assessed by plating 1 ×10^5^ cells in 6 well plates and culturing for 5 days prior to methanol fixation and Giemsa staining of surviving cells.

### Genome-wide CRISPR/Cas9 library screen

MEFs constitutively expressing Cas9 were transduced with a whole genome sgRNA library^17^ and treated with puromycin to select for a polyclonal population of sgRNA-expressing cells. Cells were treated with the BH3-mimetic ABT-737 at 250 nM (*Mcl1*^*−/−*^), 350 nM (*Mcl1*^*−/−*^*Bak*^*−/−*^ and *Mcl1*^*−/−*^*Bax*^*−/−*^) for 48 h. Surviving untreated and treated cells were harvested after 5 days, genomic DNA was extracted and enriched sgRNA were quantified by next-generation sequencing^43^. To identify genes whose sgRNAs had become significantly enriched in the surviving cell population, sgRNAs were ranked in descending order after calculating residuals to a lowest smoothed line fitted to log2-normalized counts for each sgRNA before and after selection. Minimum hypergeometric P-values were calculated from this ranked list for each gene represented in the library using an established algorithm^45^ and corrected for multiple testing.

### Subcellular fractionation, mitochondrial isolation, SDS-PAGE, blue native PAGE and immunoblotting

Cytosol and mitochondria-enriched heavy membrane fractions were fractionated following cell membrane permeabilization with 0.025% w/v digitonin as described^46^. Alternatively, mitochondria were isolated by homogenization following hypotonic cell swelling as described (Stojanovski et al, 2003).

For SDS-PAGE, lysates of whole cells or cellular fractions in reducing Laemmli sample buffer were electrophoresed through Tris-glycine gels (BioRad) and transferred to PVDF membrane.

Blue native PAGE was performed as described^47^. Proteins were transferred to PVDF, and blots were destained in 50% methanol and 25% acetic acid prior to immunoblotting.

Membranes were blocked in 5% w/v non-fat milk in TBS-T prior to immunoblotting with antibodies raised against BAK (aa23-38, #B5897 Sigma), BAK (7D10, D.C.S. Huang, Walter and Eliza Hall Institute), BAX (49F9, D.C.S. Huang, Walter and Eliza Hall Institute), BIM (3C5, L. O’Reilly, Walter and Eliza Hall Institute) and BCL-2 (3F11, L. O’Reilly, Walter and Eliza Hall Institute), cytochrome *c* (#556433, BD Biosciences), FLAG (F3165, Sigma), GAPDH (#2118, Cell Signaling Technology), HA (#11867423001, Roche), HSP70 (W. Welch, UCSF), T0MM20 (#sc-11415, Santa Cruz Biotechnologies), TIMM44 (#HPA043052, Sigma), VDAC1 (Merck, #MABN504), VDAC2 (M.T. Ryan, Monash University), VDAC3 (#55260-1-AP, Proteintech). Secondary antibodies used were horseradish peroxidase-conjugated anti-rabbit IgG, anti-mouse IgG, and anti-rat IgG (Southern Biotech).

### Generation of CRISPR/Cas9 gene-targeted mice

CRISPR/Cas9 generation of the *Vdac2/Bak/Rosa* mutant mice was performed as previously described^48, 49^. Briefly, Cas9 mRNA (20 ng/μl) and sgRNA (10 ng/μl; for sgRNA) were injected into the cytoplasm of fertilized one-cell stage embryos. 24 hours later, two-cell stage embryos were transferred into the uteri of pseudo-pregnant female mice. Viable offspring were genotyped by next-generation sequencing as previously described^43^.

### Blood and histological analysis of mice

Automated cell counts were performed on blood collected from the retro-orbital plexus into tubes containing EDTA (Sarstedt), using an Advia 2120 hematological analyzer (Siemens). Tissues were fixed in formalin, embedded, sectioned, and stained with hematoxylin and eosin.

### Mouse model of chemoresponse and tumor development

RS4;11 cells (1×10^6^) retrovirally transduced with a BFP-Luc luciferase reporter (pMSCV-IRES-Luciferase-BFP, gift from Dr. Zhen Xu) were engrafted subcutaneously into the flank of NOD/SCID/IL2Rγ mice. After 3 weeks, mice were treated for 5 days with 25 mg/kg venetoclax (ABT-199). To monitor tumor development, weekly 200 μL of 15 mg/mL D-luciferin potassium salt (Caliper Life Sciences) diluted in PBS was administered weekly by intraperitoneal injection. Fifteen min after administration of luciferin mice were anaesthetized with isoflurane inhalant and imaged using the IVIS live-imaging system (Perkin Elmer). Tumor burden was quantified by measuring the total photon flux per second emitted from the whole mouse.

To investigate tumor development, fetal liver-derived hematopoietic stem cells were harvested from mice (all on a C57BL/6 genetic background) at E14.5. Single cell suspensions were frozen prior to infection with with retrovirus expressing *c-MYC* (pMIG) in MEM supplemented with 1 mM L-glutamine, 10 mM Hepes, 1 mM sodium pyruvate, 10% (vol/vol) fetal calf serum, 50 μM β-mercatoptoethanol, and cytokines (100 ng/mL stem cell factor, 10 ng/mL IL-6, 50 ng/mL thrombopoietin, 5 ng/mL fms-related tyrosine kinase 3 ligand). Cells were then injected into lethally-irradiated (2 × 5.5 Gy, 2 h apart) recipient C57BL/6 mice. Mice were euthanized upon signs of illness (enlarged spleen or lymph nodes and weight loss). Each fetal liver was reconstituted into 6 lethally-irradiated mice with mice dying of irradiation toxicity (usually within 2 weeks) was censored from the analyses.

### Mass spectrometry and data analysis

Frozen livers were homogenized and solubilized in 1% Triton X-100. Proteins were resuspended in 6 M Urea, 100 mM DTT and 100 mM Tris-HCl pH7.0 and subjected to protein digestion using filter aided sample preparation ^50^. Peptides in MilliQ water containing 1% acetonitrile (ACN) and 1% formic acid were analyzed by nanoflow liquid chromatography tandem-mass spectrometry (LC-MS/MS) on a nanoAcquity system (Waters, Milford, MA, USA) coupled to a Q-Exactive mass spectrometer (Thermo Fisher Scientific, Bremen, Germany) through a nanoelectrospray ion source (Thermo Fisher Scientific). Peptide mixtures were loaded on a 20 mm trap column with 180 μm inner diameter (nanoAcquity UPLC 2G-V/MTrap 5 μm Symmetry C18) at 1% buffer B, and separated by reverse-phase chromatography using a 250◻mm column with 75 μm inner diameter (nanoAcquity UPLC 1.7 μm BEH130 C18) on a 120 minute linear gradient from 1% to 35% buffer B (A: 99.9% Milli-Q water, 0.1% FA; B: 99.9% ACN, 0.1% FA) at a 400 nL/min constant flow rate. The Q-Exactive was operated in a data-dependent mode, switching automatically between one full-scan and subsequent MS/MS scans of the ten most abundant peaks. The instrument was controlled using Exactive series version 2.6 and Xcalibur 3.0. Full-scans (m/z 350–1850) were acquired with a resolution of 70,000 at 200 m/z. The 10 most intense ions were sequentially isolated with a target value of 10000 ions and an isolation width of 3 m/z and fragmented using HCD with normalized collision energy of 27 and stepped collision energy of 15%. Maximum ion accumulation times were set to 50 ms for full MS scan and 150ms for MS/MS. Underfill ratio was set to 2% and dynamic exclusion was enabled and set to 60 seconds.

For the analysis of the mitochondrial BAX/BAK complexes, mitochondria from MEFs expressing FLAG-BAK, FLAG-BAX^S184L^ or untagged BAX or BAK as controls were solubilized in 1% digitonin and immunoprecipitated with anti-FLAG-coupled sepharose. Proteins were eluted with FLAG peptide in the presence of 1% digitonin, run on BN-PAGE, and stained using Sypro Ruby (Bio-Rad) and subsequently Coomassie G-250. Bands of interest were excised and subjected to in-gel reduction, alkylation, and trypsin digestion before mass spectrometry analysis as previously described^51^. Peptides from each biological replicate were run in technical triplicate, separated by reverse-phase chromatography on a 1.9 μm C18 fused silica column (I.D. 75 μm, O.D. 360 μm × 25 cm length) packed into an emitter tip (Ion Opticks, Australia), using a nano-flow HPLC (M-class, Waters). The HPLC was coupled to an Impact II UHR-QqTOF mass spectrometer (Bruker, Bremen, Germany) using a CaptiveSpray source and nanoBooster at 0.20 Bar using acetonitrile. Peptides were loaded directly onto the column at a constant flow rate of 400 nL/min with buffer A (99.9% Milli-Q water, 0.1% formic acid) and eluted with a 90 min linear gradient from 2 to 34% buffer B (99.9% acetonitrile, 0.1% formic acid).

Mass spectra were acquired in a data-dependent manner including an automatic switch between MS and MS/MS scans using a 1.5 second duty cycle and 4 Hz MS1 spectra rate followed by MS/MS scans at 8-20 Hz dependent on precursor intensity for the remainder of the cycle. MS spectra were acquired between a mass range of 200–2000 m/z. Peptide fragmentation was performed using collision-induced dissociation (CID). The raw files were analyzed using the MaxQuant software (version 1.5.2.8)^52, 53^, and extracted peaks were searched against UniProtKB/Swiss-Prot *Mus musculus* database (July 2015) containing sequences for human BAK1 and human VDAC2 and human BAX, as well as a separate reverse decoy database to empirically assess the false discovery rate (FDR) using strict trypsin specificity allowing up to 2 missed cleavages. The minimum required peptide length was set to 7 amino acids. In the main search, precursor mass tolerance was 0.006◻Da and fragment mass tolerance was 40◻ppm. The search included variable modifications of oxidation (methionine), amino-terminal acetylation, the addition of pyroglutamate (at N-termini of glutamate and glutamine) and a fixed modification of carbamidomethyl (cysteine). PSM and protein identifications were filtered using a target-decoy approach at a false discovery rate of 1%. Statistically-relevant protein expression changes between the samples were identified using a custom in-house designed pipeline as previously described^54^ where quantitation was performed at the peptide level. Probability values were corrected for multiple testing using Benjamini–Hochberg method. Cut-off lines with the function y= −log_10_(0.05)+c/(x-x_0_)^55^ were introduced to identify significantly enriched proteins. c was set to 0.2 while x_0_ was set to 1, representing proteins with a twofold (log2 protein ratios of 1 or more) or fourfold (log2 protein ratio of 2) change in protein expression, respectively.

### Bone marrow-derived hematopoietic precursor reconstitution

Bone-marrow was harvested from the femurs of *Vdac2*^*−/−*^ or age-matched wild-type control mice. Single cell suspensions were frozen at -80°C in 90% FCS/10% DMSO until reconstitution. C57BL/6 Ly5.1 mice were lethally-irradiated with 2 × 5.5 Gy 2 h apart and reconstituted with intravenous injection of hematopoietic precursors harvested from the bone marrow of *Vdac2*^*−/−*^ or age-matched C57BL/6 wild-type mice in PBS. Mice were sacrificed 8 weeks post-reconstitution and blood cell analysis was performed.

### Thymocyte and splenocyte isolation and characterization

Thymus and spleen were harvested from *Vdac2*^*−/−*^ or age-matched wild-type control mice and single cell suspensions were generated by gentle homogenization through a 100 μm sieve. Thymocytes and splenocytes were plated at 5 × 10^4^ cells per condition in 96 well flat-bottomed plates and treated for death assay. At time of analysis, thymocytes and splenocytes were harvested and stained with AnnexinV-FITC and propidium iodide for 15 mins at room temperature in AnnexinV buffer prior to analysis of viable cells (AnnexinV^−^/PI^×^) by flow cytometry.

### Platelet isolation and treatment

Platelets were isolated essentially as previously described^56^. Peripheral blood was obtained by cardiac puncture into 0.1 volume of Aster Jandl citrate-based anticoagulant (85 mM sodium citrate, 69 mM citric acid, and 20 mg/ml glucose, pH 4.6). Platelet-rich plasma was obtained by centrifugation of the murine blood diluted in buffer A (140 mM NaCl, 5 mM KCl, 12 mM trisodium citrate, 10 mM glucose, and 12.5 mM sucrose, pH 6.0) at 125 *g* for 8 min at room temperature. The supernatant was followed centrifuged at 860 *g* for 5 min and platelets were resuspended in 10 mM Hepes, 140 mM NaCl, 3 mM KCl, 0.5 mM MgCl_2_, 10 mM glucose, and 0.5 mM NaHCO_3_, pH 7.4. 40×10^6^ platelets were incubated in the presence or absence of ABT-737 for 90 min at 37°C. Viability of platelets post treatment was assessed by FITC-conjugated Annexin-V binding by flow cytometry analysis.

### Statistical analysis

Unless otherwise noted, all experiments used at least three mice per experimental group. Statistical details of the experiments including statistical tests used can be found in the Figure Legends. In the cell and animal experiments statistical significance was defined as P<0.05.

